# eGADA: enhanced Genomic Alteration Detection Algorithm, a fast genomic segmentation algorithm based on Sparse Bayesian Learning

**DOI:** 10.1101/2023.08.20.553622

**Authors:** Yu S. Huang

## Abstract

eGADA is an enhanced version of GADA, which is a fast segmentation algorithm utilizing the Sparse Bayesian Learning (or Relevance Vector Machine) technique from Tipping 2001. It can be applied to array intensity data, NGS sequencing coverage data, or any sequential data that displays characteristics of stepwise functions. Improvements by eGADA over GADA include: **a**) a customized Red-Black tree to expedite the final backward elimination step of GADA; **b**) code in C++, which is safer and better structured than C; **c**) use Boost libraries extensively to provide user-friendly help and commandline argument processing; **d**) user-friendly input and output formats; **e**) export a dynamic library eGADA.so (packaged via Boost.Python) that offers API to Python; **f**) other bug fixes/optimization. The code is published at https://github.com/polyactis/eGADA.

## Introduction

Genomic segmentation is a crucial prerequisite to detect copy number variants or alterations (CNV/CNA). The GADA algorithm ^1–3^ tackles the segmentation problem via the Sparse Bayesian Learning (also known as Relevance Vector Machine)^4^ technique to discover the minimal number of stepwise functions/wavelets (and hence the breakpoints) to describe the entire genome.

SBL used by GADA is a fast Bayesian learning algorithm. However, the backward-elimination (BE) step after SBL, which removes insignificant breakpoints, is quite slow. The BE step finds and removes the least significant breakpoint one by one. The significance of each breakpoint is established by the t-statistic comparing the coverage distribution of two flanking segments of the breakpoint. If two breakpoints have identical t-statistic, the breakpoint with the shorter length (of two flanking segments) is regarded as less significant. BE stops when two criteria have been met: **a**) t-statistics of all breakpoints are above a pre-set threshold; **b**) the number of probes/bins of each segment is above a pre-set threshold.

In terms of runtime complexity, assuming **n** breakpoints to begin with, the original GADA implementation for the BE step is of complexity **O(n**^**2**^). For a whole-genome sequencing data, **n** could be in the millions and BE becomes the most time-consuming part of GADA.

## Methods

To speed up the BE step, eGADA uses a Red-Black (RB) tree to store all segment breakpoints as nodes and then eliminate the least significant breakpoint, AKA the root of the tree, one by one. Breakpoints are sorted by their corresponding t-statistic, mentioned in the introduction. Ties are broken by the segment length of breakpoints. The segment length of a breakpoint is defined as the length of the shorter flanking segment. Red-Black tree has a time complexity of O(log(n)) for both building and querying the tree. So the time complexity of the entire BE step is improved from **O(n**^**2**^) to **O(n*log(n))**.

However, the RB tree has to be customized because removing a breakpoint involves more than just removing its node from the tree, but also merging two flanking segments into a new segment and updating the t-statistics of its two endpoints/breakpoints (and their positions in the RB tree). Here is a snippet of github eGADA/src/BaseGADA.cc .

~~~
  rbNodeType* minNodePtr = NULL;
  minNodePtr = rbTree.getMinimum();
  BreakPointKey minBPKey=minNodePtr->getKey();
  rbNodeDataType* setOfBPPtr = minNodePtr->getDataPtr();
  rbNodeDataType::iterator setOfBPIterator=(*setOfBPPtr).begin();
  //reset
  leftBreakPointPtr=NULL;
  rightBreakPointPtr=NULL;
  genomeLeftNodePtr=rbTree.nil;
  genomeRightNodePtr=rbTree.nil;

  currentMinScore = minBPKey.tscore;
  toRemoveSegmentLength = minBPKey.segmentLength;

  while (rbTree.noOfNodes()>0 && (currentMinScore<T ||
toRemoveSegmentLength<MinSegLen)){
  minBPKey = minNodePtr->getKey();
  setOfBPPtr = minNodePtr->getDataPtr();
  for (setOfBPIterator =(*setOfBPPtr).begin(); setOfBPIterator!=(*setOfBPPtr).end();
      setOfBPIterator++){
      //remove all breakpoints in this node’s data (they have same tscore and length)
      BreakPoint* minBPPtr = *setOfBPIterator; //get address of BreakPoint
      leftBreakPointPtr = minBPPtr->leftBreakPointPtr;
      rightBreakPointPtr = minBPPtr->rightBreakPointPtr;
      //update two neighboring break points.
      minBPPtr->removeItself();
      //modify genome left & right key, delete their nodes from tree and re-add them with new
key and update breakpoint info
  if (leftBreakPointPtr!=NULL && leftBreakPointPtr->nodePtr!=rbTree.nil &&
     leftBreakPointPtr->nodePtr!=NULL){
     //delete the outdated left node
    genomeLeftNodePtr = (rbNodeType*)leftBreakPointPtr->nodePtr;
    genomeLeftNodePtr->getDataPtr()->erase(leftBreakPointPtr);
    if (genomeLeftNodePtr->getDataPtr()->size()==0){
        //delete this node altogether if its vector is empty
        rbTree.deleteNode(genomeLeftNodePtr);
  }
  //new genomeLeftNodePtr that matches the new key
  genomeLeftNodePtr = rbTree.queryTree(leftBreakPointPtr->getKey());
  if (rbTree.isNULLNode(genomeLeftNodePtr)){
      //create an new node
      genomeLeftNodePtr = rbTree.insertNode(leftBreakPointPtr->getKey(),
        new rbNodeDataType() );
     }
     genomeLeftNodePtr->getDataPtr()->insert(leftBreakPointPtr);
     leftBreakPointPtr->nodePtr = genomeLeftNodePtr;
  }
  if (rightBreakPointPtr!=NULL &&
     !rbTree.isNULLNode((rbNodeType*)rightBreakPointPtr->nodePtr) &&
    rightBreakPointPtr->nodePtr!=NULL){
    //delete the outdated right node
    genomeRightNodePtr = (rbNodeType*)rightBreakPointPtr->nodePtr;
    genomeRightNodePtr->getDataPtr()->erase(rightBreakPointPtr);
    if (genomeRightNodePtr->getDataPtr()->size()==0){
        //delete this node altogether if its vector is empty
        rbTree.deleteNode(genomeRightNodePtr);
   }
   //new genomeRightNodePtr that matches the new key
   genomeRightNodePtr = rbTree.queryTree(rightBreakPointPtr->getKey());
   if (rbTree.isNULLNode(genomeRightNodePtr)){
       //create an new node
       genomeRightNodePtr = rbTree.insertNode(rightBreakPointPtr->getKey(),
       new rbNodeDataType() );
    }
    genomeRightNodePtr->getDataPtr()->insert(rightBreakPointPtr);
    rightBreakPointPtr->nodePtr = genomeRightNodePtr;
   }
  }
  (*setOfBPPtr).clear();
  //delete this minimum node after its data is all tossed out
  rbTree.deleteNode(minNodePtr);
  counter ++;
  previousRoundMinScore = minBPKey.tscore;
  previousToRemoveSegmentLength = minBPKey.segmentLength;
  if (rbTree.noOfNodes()>0){
      //get a new minimum
      minNodePtr = rbTree.getMinimum();
      minBPKey = minNodePtr->getKey();
      currentMinScore = minBPKey.tscore;
      toRemoveSegmentLength = minBPKey.segmentLength;
  }
  else{
      break;
  }
}
~~~

The Red-Black tree data structure was written in C++ template to broaden its potential applications. Here is a snippet of github.com eGADA/src/RedBlackTree.h.

~~~
template<typename keyType, typename dataType>
class RedBlackTreeNode {
public:
  keyType key;
  dataType* dataPtr;
  unsigned short color;
  /* if red=0 then the node is black */
  RedBlackTreeNode<keyType, dataType> * left;
  RedBlackTreeNode<keyType, dataType> * right;
  RedBlackTreeNode<keyType, dataType> * parent;
  RedBlackTreeNode() {
    parent = NULL;
    dataPtr = NULL;
    this->left = NULL;
    this->right = NULL;
    this->color = RED_;
  }
  /*
  * key_, data_ are references, and have to be initialized in the way above.
  */
  RedBlackTreeNode(RedBlackTreeNode<keyType, dataType>* _parent, keyType _key,
      dataType* _dataPtr) :
      parent(_parent), key(_key), dataPtr(_dataPtr) {
   this->left = NULL;
   this->right = NULL;
   this->color = RED_;
  }
  ∼RedBlackTreeNode() {
    // no memory to release?
  }
  …
  void setKey(keyType key) {
    this->key = key;
  }
  void setColor(short color) {
    this->color = color;
  }
  };
~~~

Besides using RB tree to expedite the BE step, we reorganize code into several C++ classes to better organize the source code and use Boost libraries extensively to provide user-friendly help, commandline argument processing, and user-friendly input and output formats. A dynamic Python callable library eGADA.so is also generated as part of the building process. There were also some bug fixes/optimization (reduce memory usage).

Here is a snippet to call GADA from Python after eGADA.so is built.

~~~
import eGADA
print(“### Testing the C++ eGADA.so module …\n”, flush=True)
# Pass 1 to eGADA() to enable debugging output.
# Passing 0 or no passing, i.e. eGADA.eGADA() turns off debugging.
ins = eGADA.eGADA(1)
test_vector = [1,1,1,1,0.99,0.99,1,1,0.1,0.1,0.1,0.15]
# 0.2 is alpha, 4 is min T score, 2 is min segment length.
segment_ls = ins.run(test_vector, 0.2, 4, 2)
print(f’Segmenting {test_vector} output is:\n \t {segment_ls}.\n’)
~~~

If a user encounters compiling issues, we recommend using the precompiled docker image https://hub.docker.com/repository/docker/polyactis/egada.

## Results

We ran eGADA and GADA on different inputs with identical parameters (--T 5, –alpha 0.2, –min_segment_length 0, ). Table 1 and Fig 1 are the runtime comparison results. The results confirm the theoretical time complexity analysis in the Methods section. eGADA scales log-linearly, O(n*log(n)), to n, the number of input data points, while GADA scales squarely, O(n^2^). The fraction of computing time saved will grow larger as the number of input data points increases.

**Table 1.** Runtime (in seconds) comparison between eGADA vs GADA. The raw input is https://github.com/polyactis/eGADA/blob/main/data/input.txt, which contains 80K simulated data points. The inputs are 4X, 16X, 160X of the raw data. Runtime is averaged over five repeats.

| Input | 320K data points | 1.28M data points | 6.4M data points | 12.8M data points |
| --- | --- | --- | --- | --- |
| **eGADA** | 1.88 | 8.52 | 53.60 | 102.63 |
| **GADA** | 2.85 | 13.88 | 110.74 | 326.35 |

**Fig 1.**
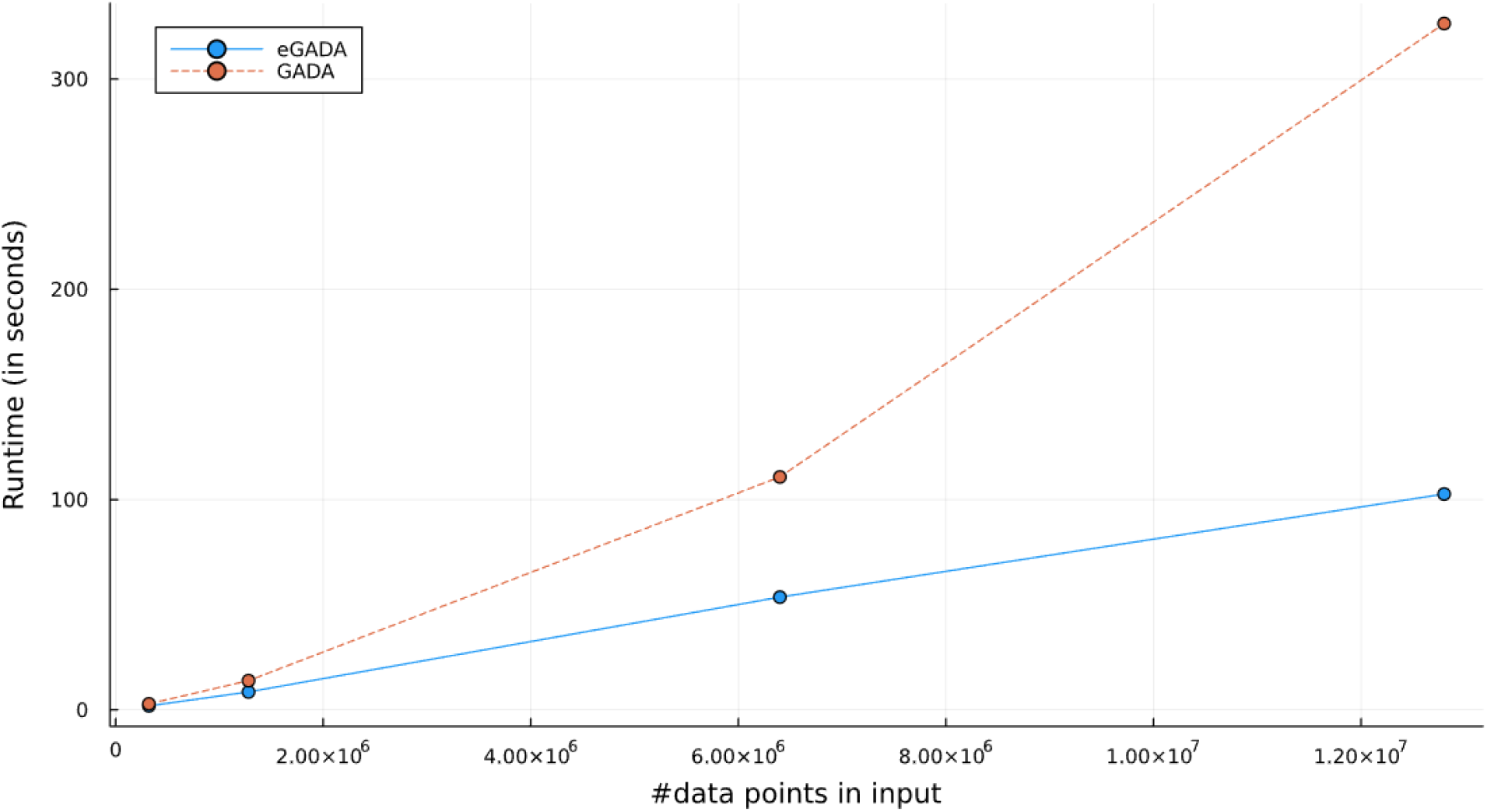
Scatterplot of Table 1. X-axis is the number of data points in each input. Y-axis is the runtime of eGADA or GADA.

We also compared eGADA with BIC-seq2 ^6^, in segmenting the genomic data (each input data point is normalized coverage of a 500bp bin) of simulated and TCGA samples, and found eGADA can produce similar segmentation results (data not shown) while being much faster. We included it as part of Accucopy, a tumor-purity and CNA inference software^5^.

## Acknowledgement

Most of the development of eGADA took place while the author was a graduate student at USC, befriending Dr. Pique-Regi, the author of GADA, during research collaboration. The code was polished over the years. Thank Dr. Pique-Regi for several discussions about the algorithm. Thank Dr. Xinping Fan for fixing a bug in the RB C++ template code.

